# Refining modules to determine functionally significant clusters in molecular networks

**DOI:** 10.1101/756064

**Authors:** Rama Kaalia, Jagath C. Rajapakse

## Abstract

Module detection algorithms relying on modularity maximization suffer from an inherent resolution limit that hinders detection of small topological modules, especially in molecular networks where most biological processes are believed to form small and compact communities. We propose a novel modular refinement approach that helps finding functionally significant modules of molecular networks. The module refinement algorithm improves the quality of topological modules in protein-protein interaction networks by finding biologically functionally significant modules. The algorithm is based on the fact that functional modules in biology do not necessarily represent those corresponding to maximum modularity. Larger modules corresponding to maximal modularity are incrementally re-modularized again under specific constraints so that smaller yet topologically and biologically valid modules are recovered. We show improvement in quality and functional coverage of modules using experiments on synthetic and real protein-protein interaction networks. Results were also compared with six existing methods available for clustering biological networks. In conclusion, the proposed algorithm finds smaller but functionally relevant modules that are undetected by classical quality maximization approaches for modular detection. The refinement procedure helps to detect more functionally enriched modules in protein-protein interaction networks, which are also more coherent with functionally characterised gene sets.

## Introduction

Module detection in molecular networks has helped in identifying protein complexes, biological pathways, and functional and disease modules^1–4^. Many algorithms have been introduced to obtain biologically significant modules of genes, proteins and metabolites^5–13^. These algorithms use topological properties of networks to cluster nodes into modules; a module being defined as a subgraph of nodes, having more dense connections among themselves than with the rest of the network. Examples of such few state-of-art algorithms are Newman-Girvan^5,9^, Louvain^7^, InfoMap^14^, Spinglass^15^, Random Walks^16^ and Markov Cluster algorithm^17^. Research advances in protein function associations^18–20^ have also led to development of few clustering approaches designed specifically for protein interaction networks to detect protein complexes and functional modules such as Molecular Complex Detection (MCODE)^10^, DPCLUS^11,13^, Clustering with Overlapping Neighbourhood Expansion (ClusterONE)^12^ and ModuleDiscoverer^21^.

Modularity^5^ was introduced by Newman *et. al*. and has pioneered the works in identifying modules in physical and biological sciences. Modularity based module detection is based on rearranging nodes in modules to maximize the modularity of the resulting partitioning^5,6,9,22^. These algorithms have shown good performances in many biological applications^23^ but suffer from a resolution limit as they fail to detect small modules^24^. The other global quality functions mathematically similar to modularity, where the quality of a partition is given by the sum of the qualities of the individual modules, also have a resolution limit because of the trade-off between the number of modules and the quality of each module. This combined with the fact that biologically significant functional modules are smaller than topological modules detected using such algorithms requires an approach that is able to detect smaller but functionally valid modules specific to biological networks. A few algorithms have been introduced to address this problem like Spinglass^25^, Random Walks^16^ and Asymptotical Surprise^26^. Despite the modularity based algorithms outperforming in modularization^23^ and the modularity being the only widely accepted metric available for evaluating communities, there is limited research done till date to address the need of methods for finding functionally significant smaller clusters especially in the applications to molecular networks. The existing approaches for finding modules are ineffective for molecular networks as functional clusters do not necessarily correspond to the partitions of genes obtained by maximizing the modularity^27^. Biological complexity results in many interactions within (intra) and with other (inter) functional modules, resulting in functional clusters with lower modularity values which is further compounded by incomplete nature of these networks.

In the present study, we propose a new module refinement algorithm that refines the modules obtained from any modularity based community detection method. Our module refinement method lowers the quality (modularity) of modularization within the sub-optimal zone of modularity and incrementally re-invent new modules from the larger modules having possible sub-modules. All the while specific constraints are maintained so that only small modules that fit into the module definition under the topological constraints are refined. We show the improvement in topological and functional quality of modules detected after applying our refinement method by conducting various experiments on benchmark synthetic networks and human protein-protein interaction networks along with the comparison with six existing module detection algorithms.

## Results

First set of experiments were performed on benchmark synthetic networks and real human protein-protein interaction networks (PPIN). For real networks, the modules predicted using our module refinement algorithm (see Methods for details) were compared to the two modularity optimizing algorithms Louvain^7^ (L) and Clauset-Newman-Moore Greedy^6^ (G) algorithm, one node property based algorithm Label Propagation^28^ (LP), one resolution limit free algorithm Asymptotical Surprise^29^ (ASY) and two seed based clustering algorithms MCODE^10^ and DPCLUS^13^. The algorithms were developed in Python^30^ (version 3.6.1) and import modules from python packages NetworkX^31^ (version 2.2) and Community^7,32^ (version 0.11).

### Experiments with Synthetic Networks

#### Benchmark Synthetic Networks

Molecular networks such as PPIN show heterogeneity in the sizes and degree distributions of modules. We generated benchmark networks by using the LFR algorithm (Lancichinetti, Fortunato & Radicchi)^33^. It uses power-law distributions of node degrees and community sizes to generate real world like synthetic networks. PPIN like most real networks are far from complete and lack ground truth of modules to compare with, so LFR benchmark networks were used to quantitatively evaluate the performance of our method. LFR networks are especially suited for evaluating our algorithm as (1) they can use power-law distributions of node degrees and community sizes to generate real world like networks; and (2) they can use different values of mixing parameter to define the fraction of inter and intra-module edges and thus represent various degrees of incompleteness and interconnectedness in real biological networks. For present study, we randomly generated 50 benchmark networks for each set of parameters (Table 1). The parameters were selected using prior knowledge of PPIN and hence mimic the characteristics of molecular networks.

**Table 1.**
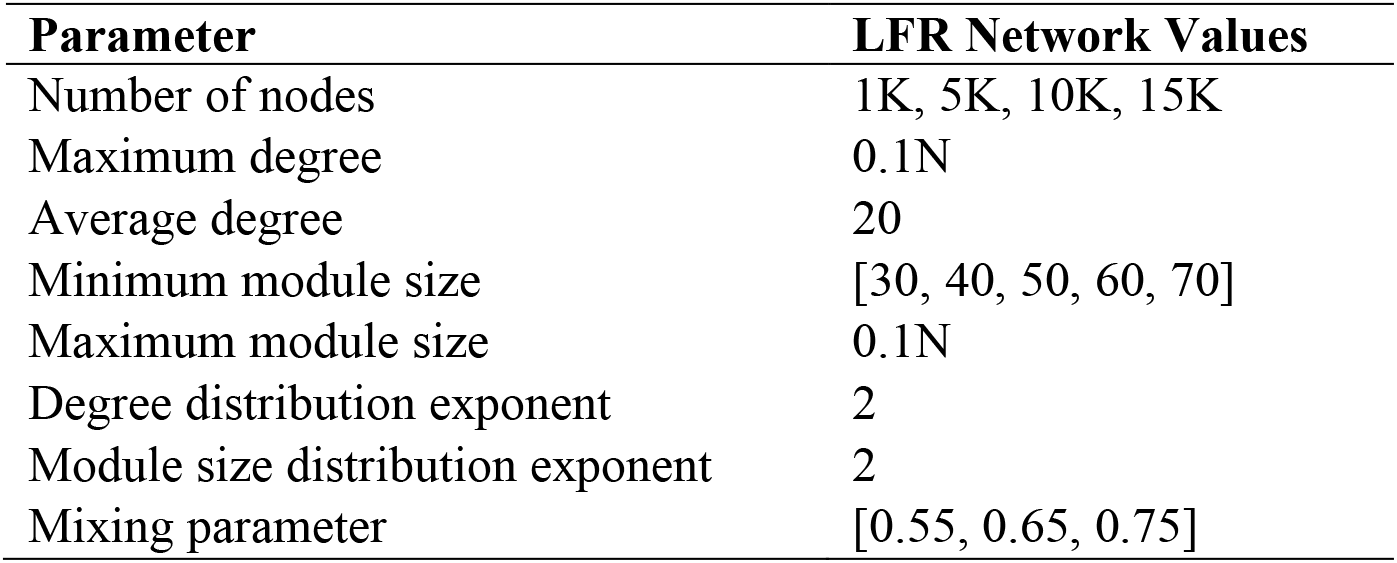
Parameters used to generate LFR benchmark networks

#### Performance Evaluation

Benchmark networks were modularized using only Louvain at resolution parameter, γ=1 and with module refinement method. We evaluated the performance of our method using Normalised Mutual Information (NMI) values (to measure accuracy w.r.t. ground truths), the modularity, size of the modules and computational complexity (time) for benchmark networks of different sizes (N=1000, 5000, 10000, 15000) at different values of mixing parameter. Figure 1 shows the mean values of performance metrics for Louvain and Louvain with module refinement over 50 benchmark networks. The performance metrics for N=1000 were observed to be quite low indicating the dependency of modularization on number of nodes in a network. As observed in the Figure 1, after applying refinement in module detection, we observed accuracies of 76.3%, 76.6% and 81% in identifying modules as compared to 72.1%, 70.5% and 75.2% accuracies of Louvain algorithm alone for networks of sizes 5K, 10K and 15K, respectively. The accuracy of module detection decreased with increase in mixing parameter. After refinement, modules had sub-optimal modularity values but more accurate with high NMI values. More importantly, refined modules are of smaller size of less than 200 as compared to Louvain alone that produced large modules as big as of size 1200 which are unlikely to have biological relevance. Because of the incremental modularization step, refinement method takes thrice as much time as taken by just Louvain. Overall, with refinement modularization is found to be more accurate.

**Figure 1.**
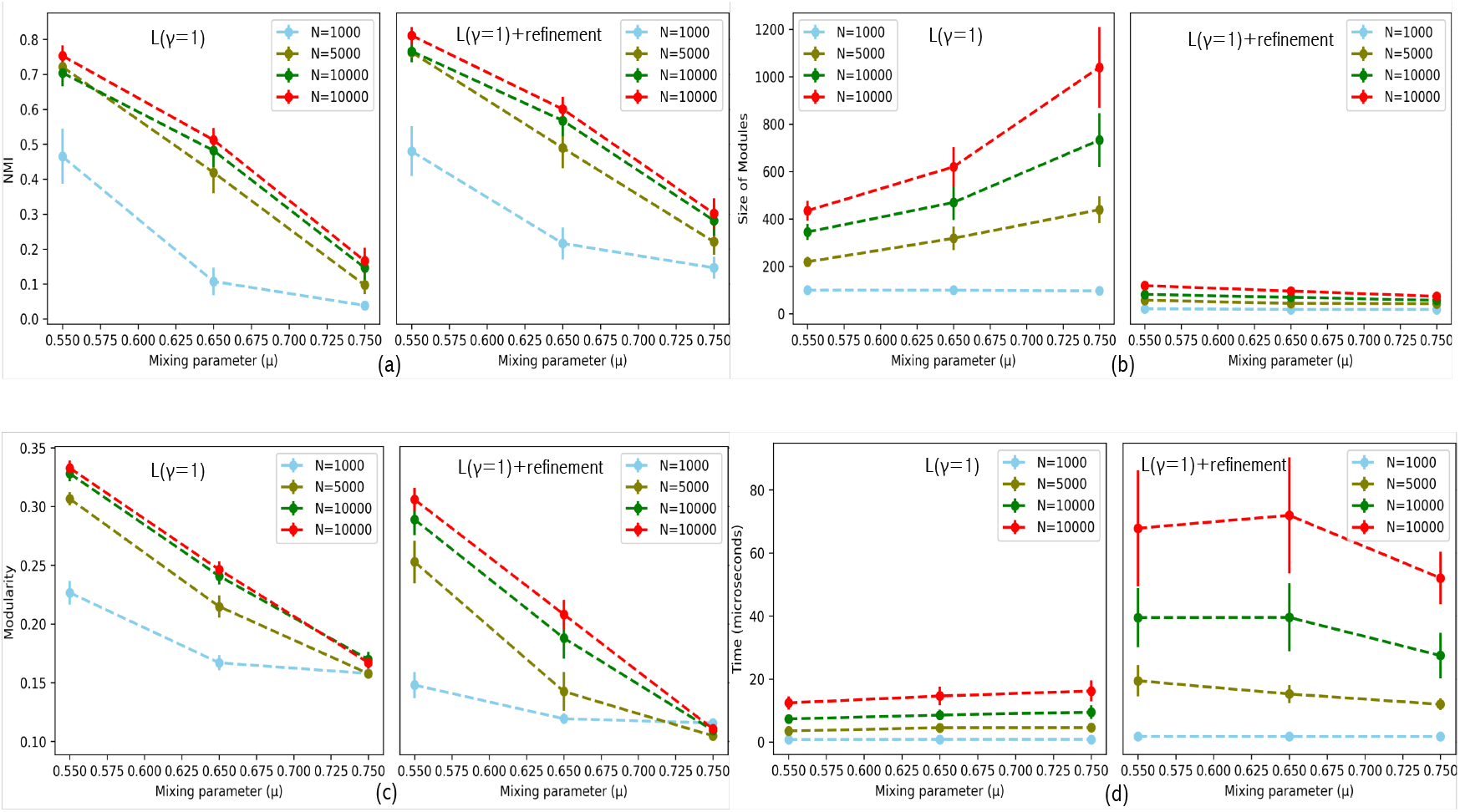
Performance comparison of just Louvain (γ=1) (left) and Louvain (γ=1) with refinement (right) on LFR benchmark networks. (a) NMI values, (b) Modularity (*Q*), (c) Size of modules, and (d) Computing time (in *µ*s). The x-axis represents the mixing parameters, y-axis represents the mean values of the performance metric, and error bars show standard deviations.

### Experiments with Protein-Protein Interaction Networks

#### Protein-Protein Interaction Networks (PPIN)

A comprehensive set of Human PPIN was constructed using protein-protein interactions retrieved from STRING^18^, HPRD^34^, BIOGRID^35^ and IMEx consortium databases^36^ (DIP, IntAct, MINT, HPIDB, UniProt etc). High confidence interactions from STRING and IMEx databases along with curated experimental interactions from HPRD and BIOGRID were combined to create a PPIN dataset that contains 78705 interactions and 12022 nodes with average (maximum) degree of 13 (496) (see Additional file 1 for details).

#### Evaluation on Human PPIN

Modules were detected in PPIN by optimizing modularity using Louvain (L) and Clauset-Newman-Moore Greedy (G) algorithms. The observed modules were re-modularized into smaller modules using the refinement algorithm developed in this study. It shows that our refinement method can be implemented on a partition obtained from any modularity based algorithm. Performances of modularization were also compared with four non-modularity algorithms such as Label Propagation (LP), Asymptotical Surprise (ASY), MCODE and DPCLUS. We claim that our refinement method also addresses the problem of resolution limit which is substantiated by the comparison provided by ASY method.

#### Refining Modules at different resolution parameters (γ)

Modularity based optimization suffers from a resolution limit. To address this limitation, a resolution parameter, γ was introduced by Reichardt and Bornholdt^15^ in order to detect finer modules such that original modularity^5^ is kept at γ =1 (eq(1)). We first show that clustering PPIN using just modularity optimization with Louvain algorithm at different values of γ is less effective than our refinement procedure in addressing the resolution limit problem in PPIN.

Performance at different γ values is measured using mean values of pairwise NMI, modularity, number and size of modules over 25 iterations of the Louvain. As seen in Figure 2, For γ values from 0 to 1.5, stability of module detection is very low, modularity values are high but that is because of few very large modules detected in these partitions. From γ values of 2 onwards, high values of pairwise NMI over iterations show stability in module detection using our method as compared to those from Louvain alone. Also, refinement procedure results in more and smaller modules from PPIN but with lesser modularity values as expected.

**Figure 2.**
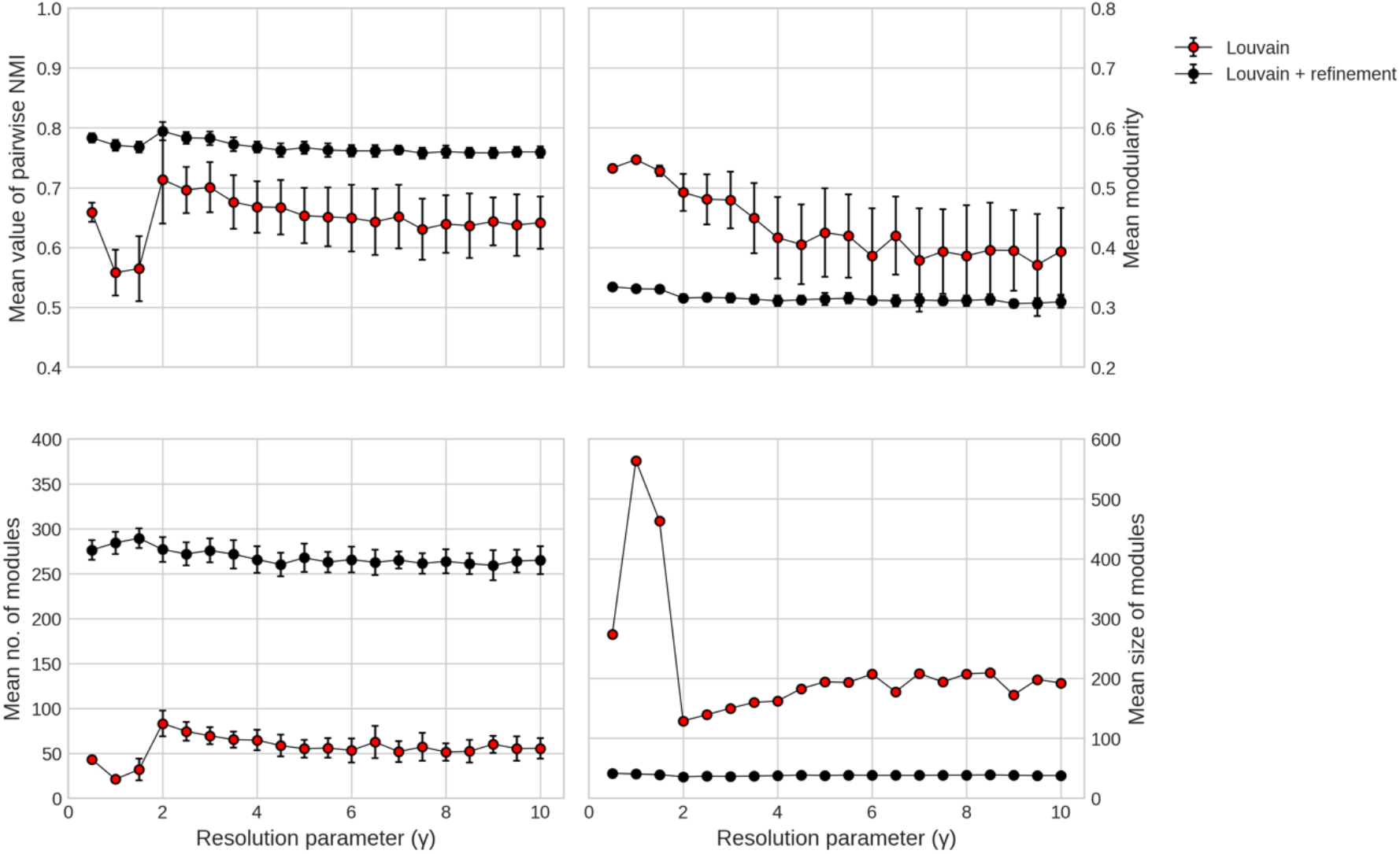
Performance of just Louvain and Louvain with refinement at different γ values. Red circles depict partition from just Louvain and black circles depict partition from Louvain with refinement with error bars for NMI, modularity and number of modules.

#### Selecting best resolution parameter

To select the best clustering of PPIN, a value of resolution parameter is selected for modularity optimization using Louvain that corresponds to high stability and significance of modules. Maximizing modularity with Louvain has a stochastic element and thus stability is measured using mean of pairwise NMI across 25 iterations of algorithm run. Significance of modules is calculated by adapting the approach from^37^ where we compared the modularity values from 25 iterations of PPIN with 100 random networks generated with same degree sequence as the PPIN. Figure 3 shows that modularity values of partition from Louvain are significantly larger for γ in range of 1.5 to 10. But as observed in Figure 2, most stable, smaller and more number of modules are given by γ = 2. For further comparisons and evaluation, clustering of PPIN performed with Louvain at this best γ value is considered and the refinement procedure developed in this study is applied to these clusters. We will refer to original modularity optimization with just Louvain at γ = 2 as L(γ = 2) and with our refinement procedure as L(γ = 2) with refinement.

**Figure 3.**
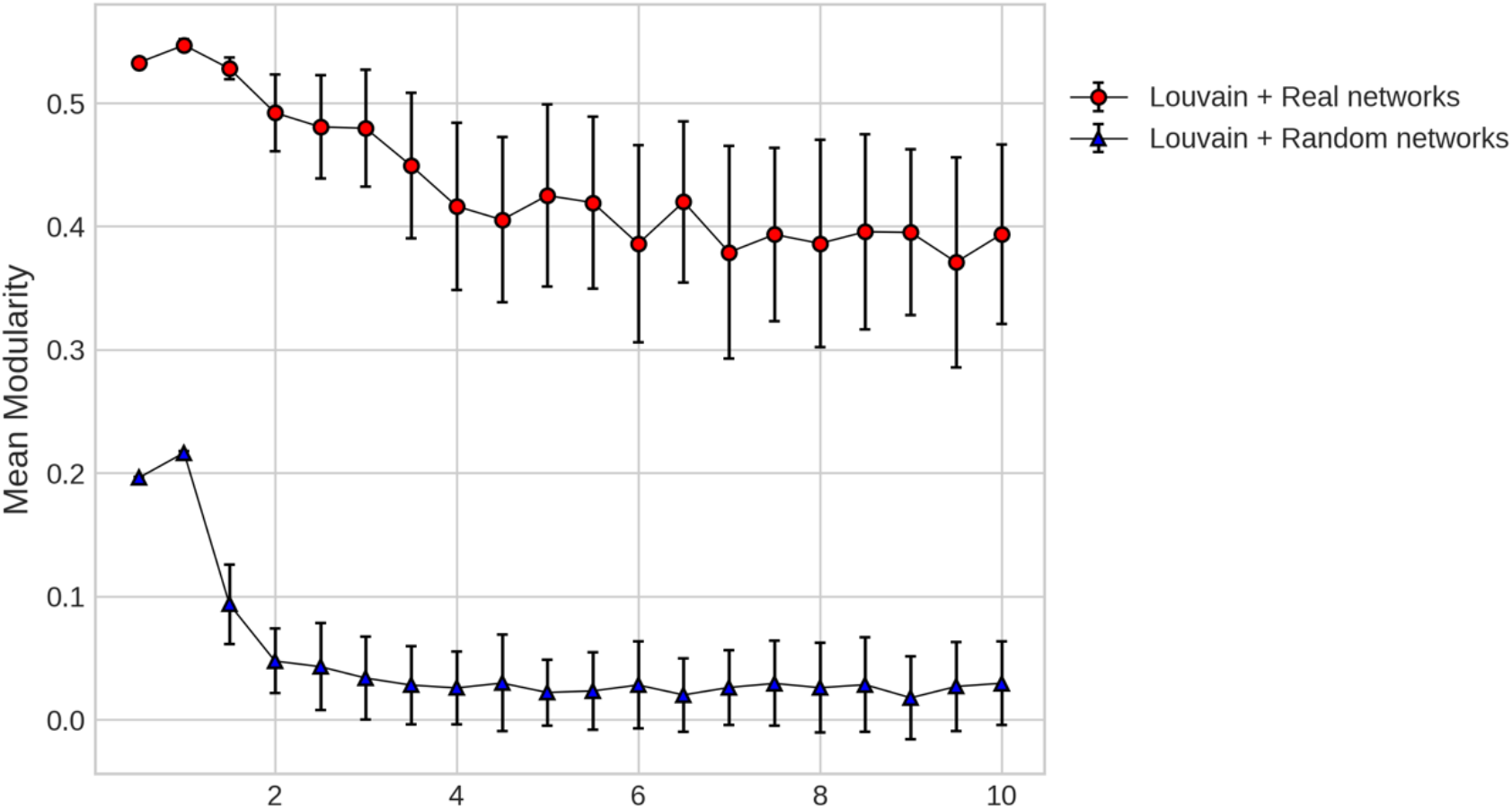
Mean modularity values for partition obtained from Louvain at different values of resolution parameter. Red shows real protein-protein interaction networks and Blue shows random networks of same degree sequence.

#### Effect of module refinement

Sizes of modules obtained from only Louvain at γ =1 and 2, and after refinement were studied to see the effectiveness of the refinement algorithm on the initial modularization. As expected, modules after refinement are smaller for both γ values. Refinable modules from L(γ) (that can be modularized further) and non-refinable (those that do not contain sub-modules) modules from L(γ) are shown in Figure 4. The refinable modules are processed using refinement algorithm to produce new refined modules. Figure 5 shows size distribution of these three types of modules. See Additional file 1 Figure S1 and S2 for size distribution of modules for L(γ=2). For L(γ=1), 58% of 25 modules from initial modularization were refinable and resulted in 305 modules in final modularization after refinement. Whereas for L(γ=2), 13% of refinable modules out of 96 resulted in 282 modules in final modularization.

**Figure 4.**
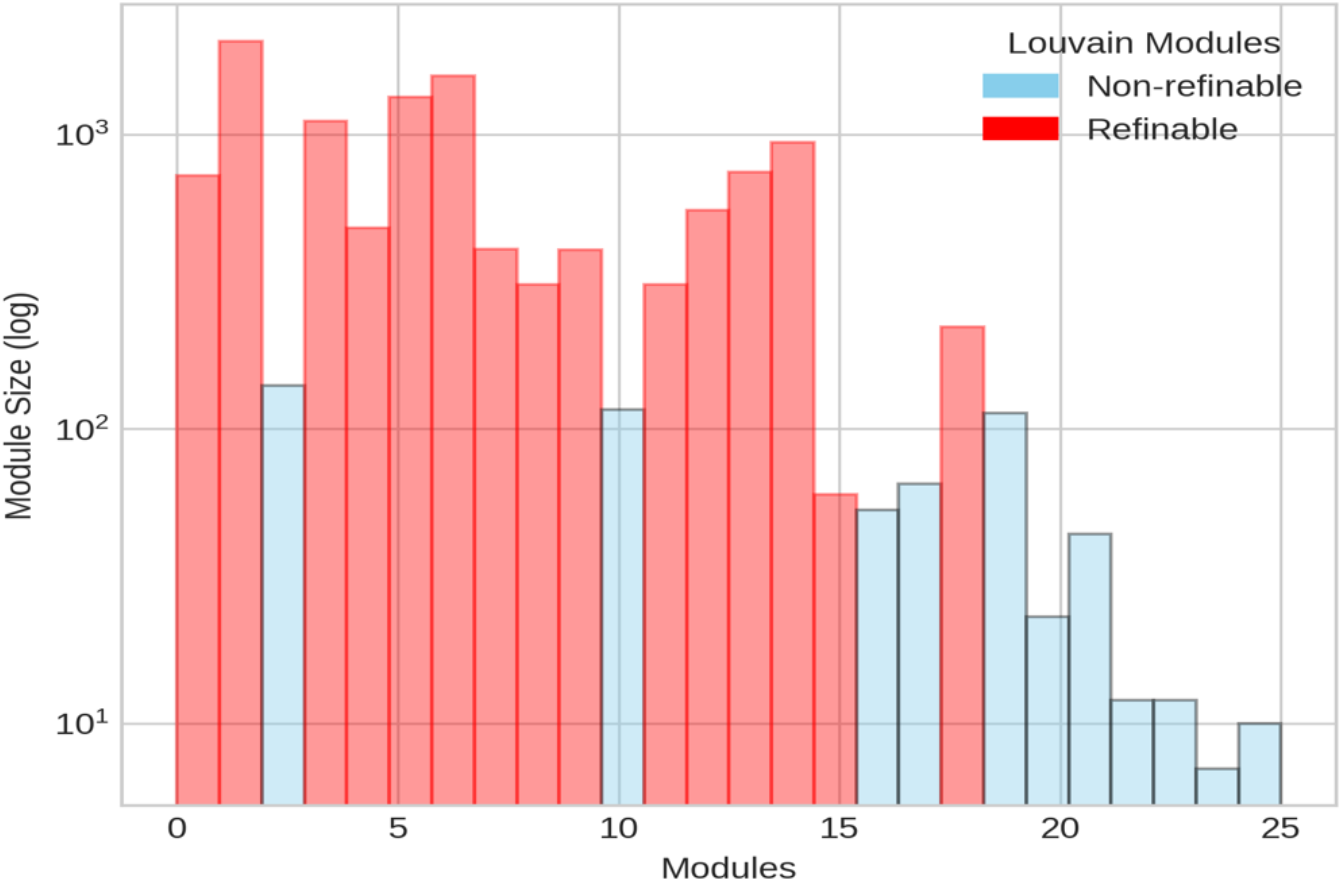
Size distribution of refinable and non-refinable modules obtained from Louvain based modularity optimization, L(γ=1).

**Figure 5.**
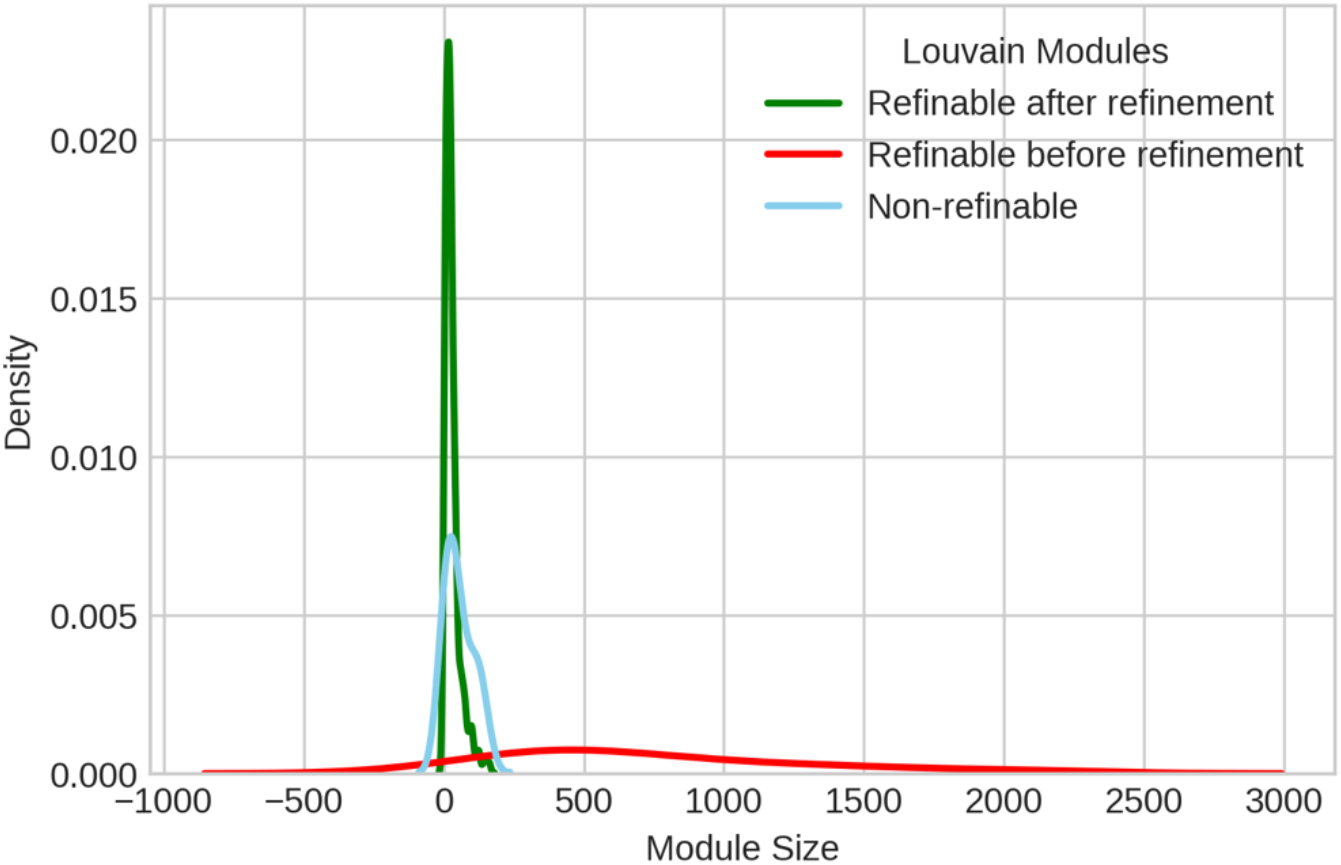
Module size distribution of refinable (from L(γ=1)), non-refinable (from L(γ=1)) and refined (L(γ=1) +refinement) modules.

#### Comparison with other Algorithms

Performance of the refinement method was tested with not only just Louvain (L), but with one more modularity based algorithm (Clauset-Newman-Moore greedy optimization (G)) along with a node property based algorithm (Label propagation (LP)), a resolution limit free algorithm (asymptotic surprise (ASY)) and two PPIN clustering algorithms that use weighted vertices as seeds for initial clusters. Figure 6 represents complementary cumulative distribution function for size of modules from different algorithms. ASY modularizes a network by optimizing a probability based quality function and addresses the resolution limit problem by estimating smaller modules. L, G and LP all detect many large modules of size more than 500. MCODE and DPCLUS are designed specifically to detect small and dense protein complexes. MCODE modules are observed to be of similar size distribution to our refinement algorithm while DPCLUS reported 88% modules with no more than three protein nodes and only 4% modules with more than ten protein nodes (meso-modules). It is clear that if large modules obtained from G and L are processed using our refinement step, smaller modules comparable to resolution limit free algorithms are obtained that are more likely to be closer to functional modules.

**Figure 6.**
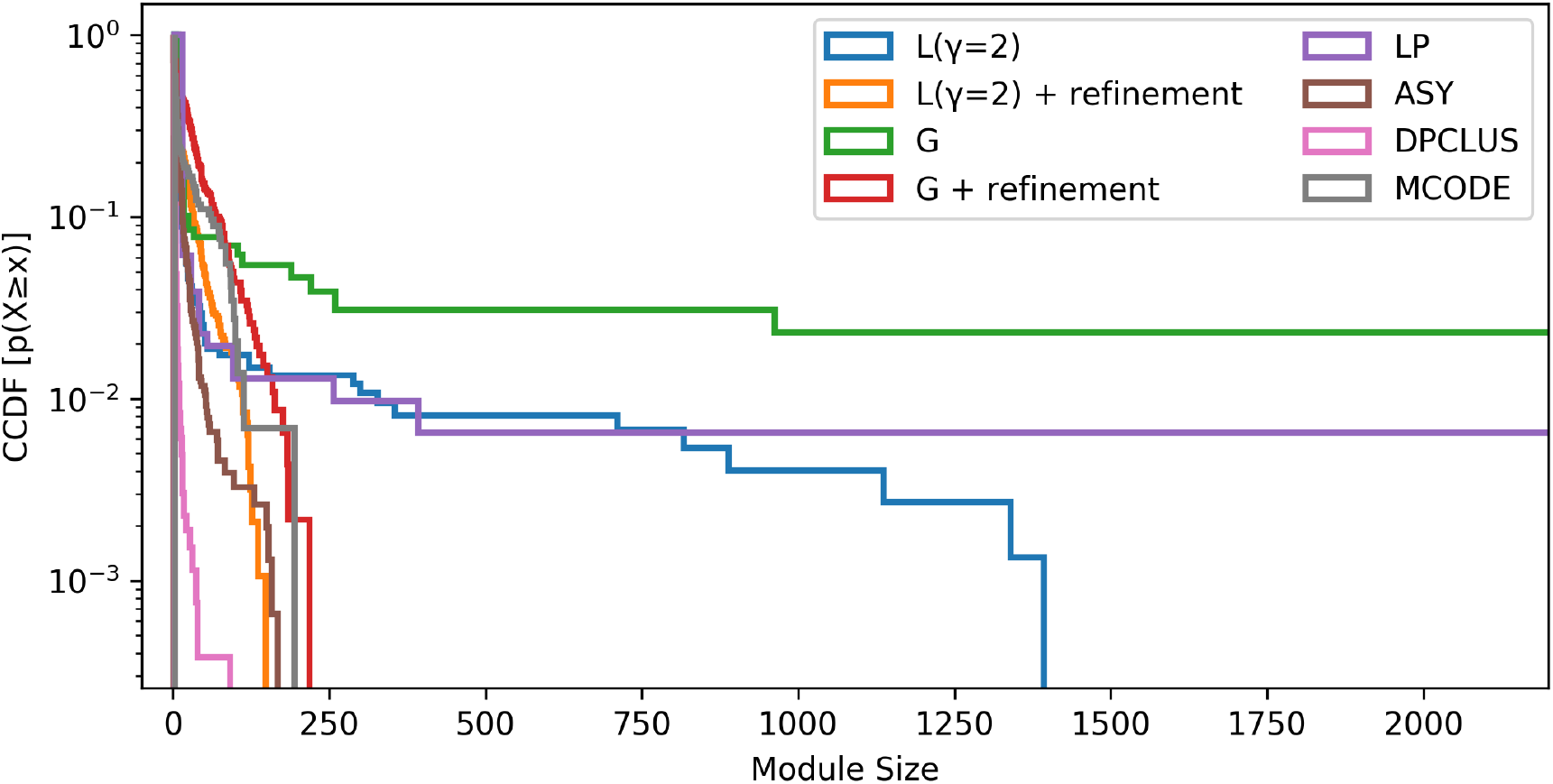
Complementary cumulative distribution function for size of modules from different algorithms: Louvain (L), Clauset-Newman-Moore greedy optimization (G), Label propagation (LP), Asymptotic surprise (ASY), MCODE and DPCLUS.

Structural quality of modules was calculated using a composite performance score that consists of modularity and partition density (see Methods for details). It measures the topological quality of the modules in a partition. Partition density represents edge density of the resulting clusters. As observed in Figure 7, modularity and partition density are higher for ASY, MCODE and DPCLUS. Modularity optimization methods (L and G) achieve high partition density only with refinement. Overall performance and partition density is found to be better for modules obtained from DPCLUS, ASY and Louvain with refinement algorithms.

**Figure 7.**
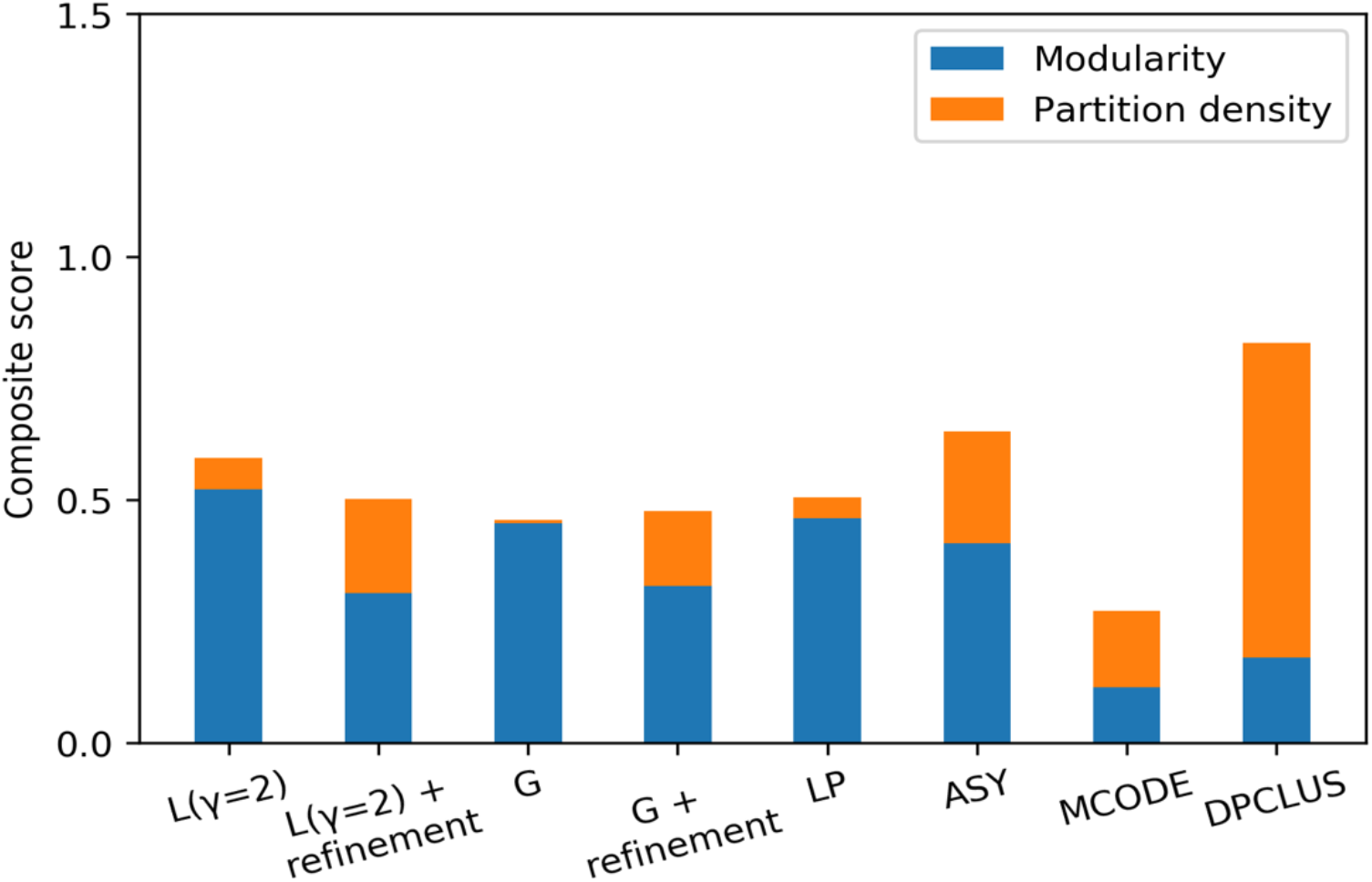
Composite performance of modules given by different algorithms-Louvain (L), Clauset-Newman-Moore Greedy optimization (G), Label propagation (LP), Asymptotic surprise (ASY), MCODE and DPCLUS.

#### Functional Enrichment of Modules

In order to evaluate the biological relevance of the topological modules detected from PPIN we determine GO-MF terms that are significantly enriched in the meso-modules (size>10). Significantly enriched functions were evaluated at *p*-value=0.05 by using the GOstats^38^ package. Figure 8 sheds light on how the refined modules are highly enriched in molecular functions despite their smaller sizes as compared to the larger modules detected by the Louvain algorithm. The average fractions of functionally important genes in a module, calculated with respect to top ten enriched functions, were much higher for refined modules than those detected by the modularity optimization algorithms (L,G) and other algorithms (LP, ASY). Though asymptotical surprise produced smaller modules with high topological quality, their functional relevance is quite low as compared to refined modules from our method. MCODE modules showed comparable enrichment of functional genes and DPCLUS showed twice as much enrichment. On closer inspection, we found that only 33 out of 774 DPCLUS clusters are meso-modules and contributed to functional enrichment whereas Louvain with refinement produced 283 enriched clusters out of 678 refined modules. Therefore, apart from being topologically valid and smaller in size, the refinement produces more modules that are clustered with similar biological functions.

**Figure 8.**
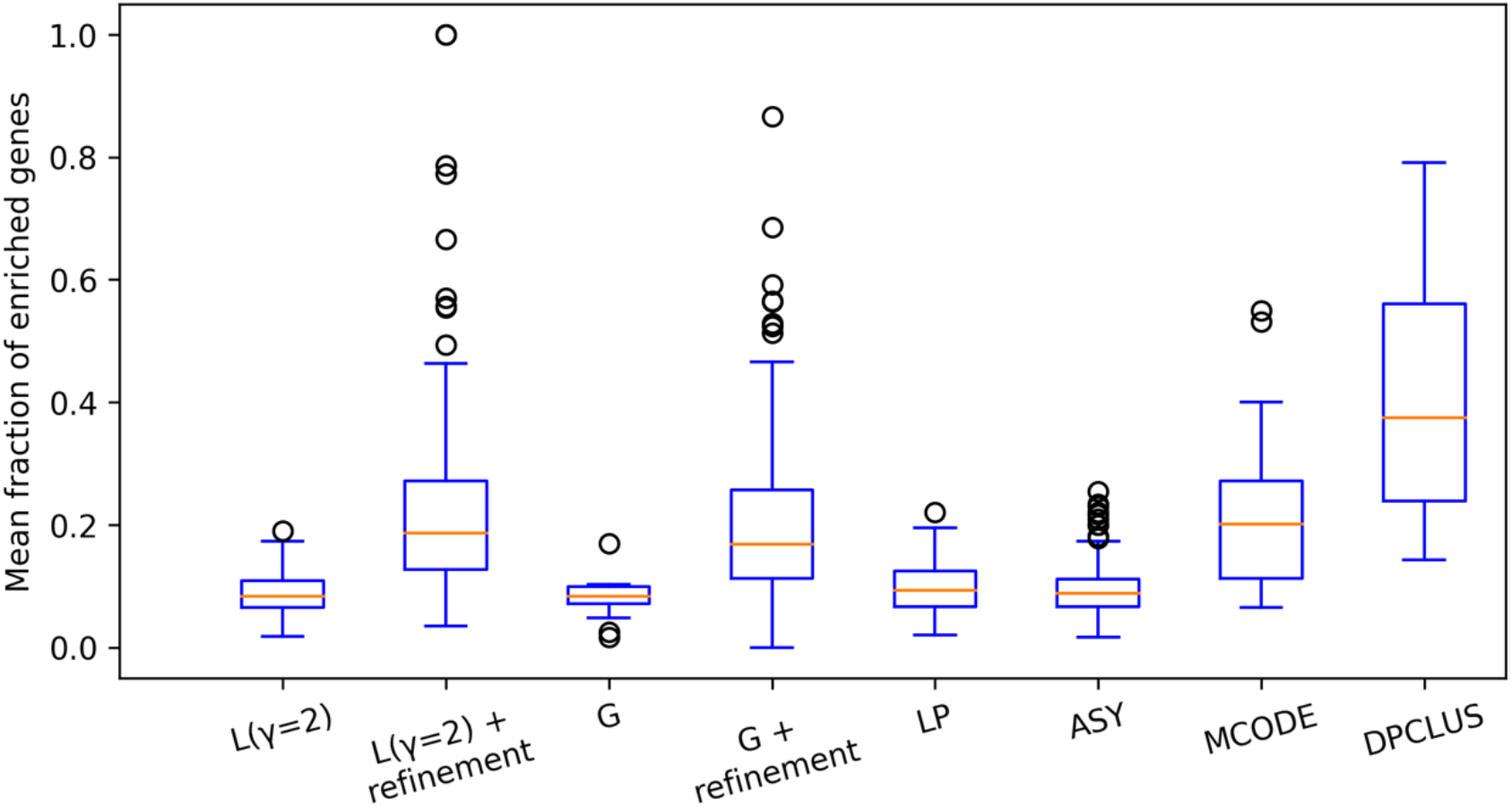
Mean percentage of functionally significant genes in modules from different algorithms: Louvain (L), Clauset-Newman-Moore Greedy optimization (G), Label propagation (LP), Asymptotic surprise (ASY), MCODE and DPCLUS. Only meso-modules (with size>10) are considered.

#### Functional Validation using Known Gene Sets

To experimentally validate the modularization algorithms, we employ 186 KEGG^39^ pathways and 50 Hallmark gene sets (MSigDB^40^) that are well characterised by their biological processes. We estimated the overlap between these biological processes and the refined modules from L(γ = 2) by using the functional coverage (see Methods section). The functional coverage is given by average overlap between the known functions and predicted clusters. Modules from L(γ = 2) showed functional coverage of 3.9 and 4 % for Hallmark gene sets and KEGG pathways respectively. Applying refinement method on these modules drastically increased the coverage to 8.4 and 15.6 % respectively. Thus, the refinement of modules from human PPIN resulted in topological modules that are more closer to known functionally characterised gene sets with more than two fold increase in functional overlap compared to those detected by using just modularity optimization algorithms. Note that overall low coverage values cannot be avoided due to the incompleteness of functionally characterised gene sets.

#### Functional Significance of Refined Modules: a case study

In this section, we attempt to shed light on the biological properties of the refined modules realized by our algorithm. An in-depth analysis of individual modules confirmed that after refinement, the observed modules become more specific to few biological functions. For example, we studied a refinable module, *X* from Louvain at γ=2 in detail (see Additional File 1 for gene list). *X* can be considered as a supermodule that is re-modularized into four sub-modules *A*, *B*, *C* and *D*. Figure 9 shows the different functional categories of the supermodule and its submodules after refinement using PANTHER^41^ classification system for protein functions and classes.

**Figure 9.**
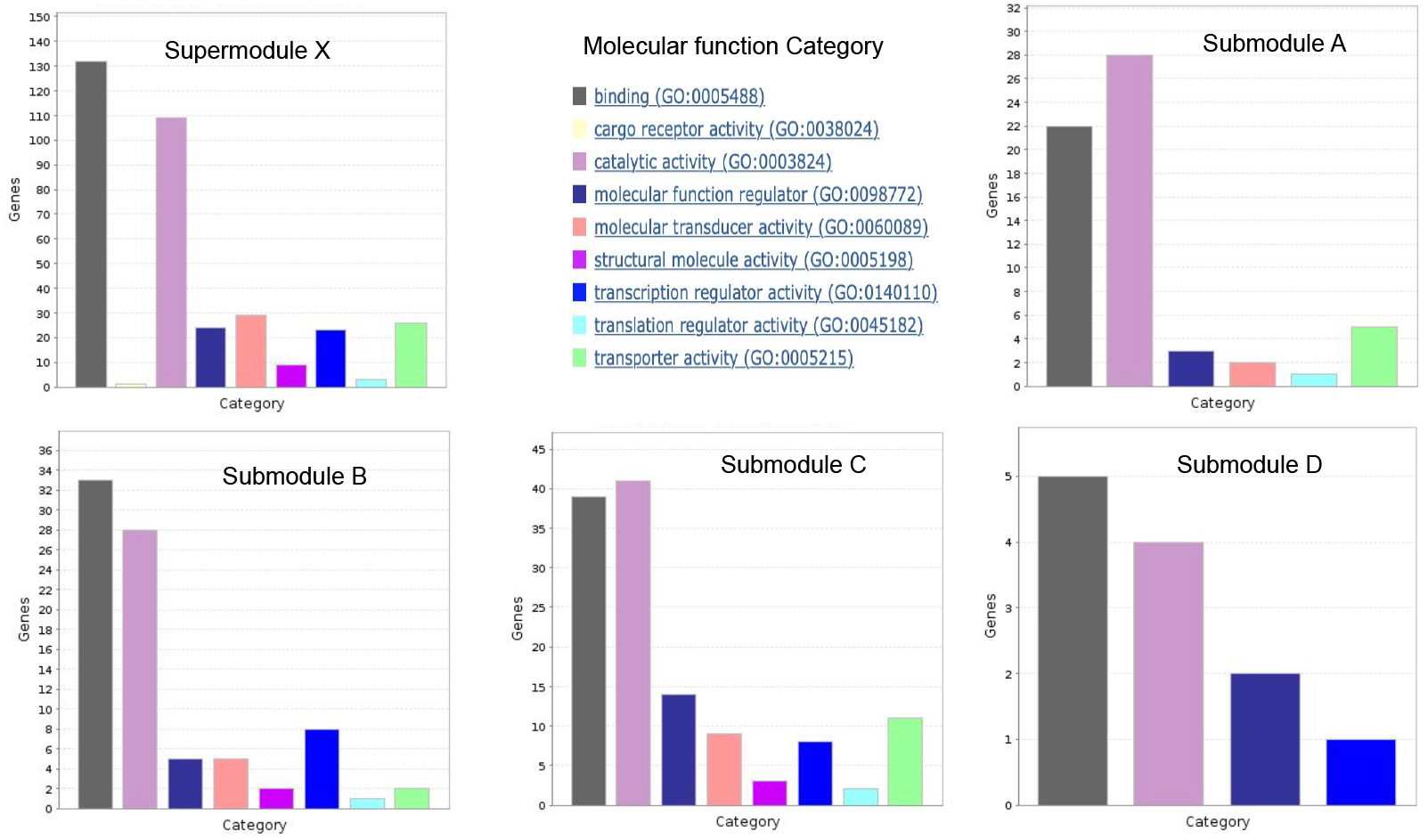
Functional categories of genes in the supermodule (*X*) and its refined submodules (*A*, *B*, *C*, *D*). The figures for functional classification are generated using web application of PANTHER classification system^41^.

The supermodule *X* is found to be rich in following protein functions:

1. Binding and catalysis: 37% and 31% of proteins in the supermodule *X* bind with other biomolecules and catalyze reactions, respectively. They include non-receptor serine/threonine protein kinases (e.g., CAMKV, MAPK8, PRKDC, ATR), proteases (e.g., PRSS1, DPP3, PSEN2), RNA polymerase, DNA polymerase and calcium binding proteins (e.g., CAB39, PLCG2).
2. Signal transducers: 8% of proteins are involved in signaling processes such as G-protein coupled receptors (e.g., MC3R, GPR6, AVPR2, SSTR5, GALR3, HTR2B, SSTR3, HTR1D) and cytokine receptors (e.g., IL20RA and IL20RB)
3. Transporter: 7% of proteins (e.g., SLC6A12, FXYD3, SLC9A3, SLCO1A2, FXYD6, SLC1A5, TRPM6, SLC10A2, SLC5A7, KCNQ1, KCND2) are involved in molecular transport.
4. Regulators: 14% of proteins in the module are responsible for regulation, activation and inhibition of various biomolecules. They include functional regulators such as G-protein modulator (e.g., RGS 16, MYO5C, MYO15A, EPS15, ERC2, EV15L, RGS16), kinase activators (e.g., DBF4B, CCNJL) and protease inhibitors (e.g., WFDC2, ITIH3, BIRC2); transcriptional regulators such as transcription factors (e.g., SIX1, PLAG1, SOX3, MAF, STAT5A, EGR4, TAF10, MAFG, RUNX1T1, TBP, SMYD1, FOXI1, SIX2, C1D, MYCL) and translational regulators like RNA binding proteins (e.g., DAZL, DAZ4) and translation initiation factor (e.g., EIF2S2).
5. Structural molecules: 3% of proteins like tubulin (e.g., TUBA1A, TUBB2B), collagen (e.g., COL18A1), actin binding motor proteins (e.g., MYO5C, MYO15A), ribosomal proteins (e.g., GRSF1, RPL17-C18orf32) belong to class of structural proteins.

Submodule *A* is found to be mainly comprised of heat shock proteins such as HSPA1A, HSPA1B, HSPA1L, HSPBP1, HSPA2, HSPA5, HSPA6, and HSPA8, which stabilize proteins when exposed to heat; and transmembrane transporters like voltage-gated potassium channel (e.g., KCNE2, KCNH2) proteins, calcium channel (e.g., CACNA1E) proteins and cotransporters (e.g., SLC5A1 and SLC12A3). Submodule *B* is enriched more in chaperone proteins (e.g., HSP90AB1, HSP90AA1, CDC37L1, CDC37, PGGES3, FKBPL) responsible for correct folding and transport of unfolded proteins; kinases that regulate multiple cellular signalling processes (e.g., CDK15, CDK11A, MAP3K9, E1F2AK1, IP6K2) and transcription factors (e.g., SM4D2, SM4D3, ELK4, MAFK, ELFS, HOXD4). Submodule *C* is highly enriched in sets of regulator and signalling proteins such as serine/threonine kinases (e.g., CDK11B, CDK14, CDK16, CDK17, SIK3, SK1, PAK4, DYRK1A), G-protein coupled receptors (GPCR) (e.g., ADRA2B, ADRA2C) and G-protein modulators (e.g., TBC1D4, MPR1P, RAP1GAP2, RIN1, TBC1D4, RALGPS2, GAPVD1, RASSF1) that activate and regulate many signalling molecules; it is also composed of a number of transmembrane transporters and voltage-gated ion channels (e.g., CACNB2, KCNK15, KCNK9, KCNK3). Module *D* is a relatively smaller module with 12 functional mappings to its gene nodes including G-protein modulators and casein kinases that are involved in signalling process and endocytosis involving vesicle formation from membrane, respectively. In the refined submodules, proteins from same family and functions are found to be clustered together.

## Discussion

The refinement algorithm developed in this study is based on re-modularizing the refinable modules generated by a modularity optimizing algorithm. For the present study, we chose two state-of-art modularity optimizing algorithms-Louvain and Clauset-Newman-Moore Greedy optimization because of their recent popularity, especially in applications of molecular networks. One node property based algorithm (Label Propagation) and one resolution limit free algorithm (Asymptotical Surprise) were also selected for comparison. For a comprehensive evaluation, modularization with two seed based algorithms specific to protein interaction networks (MCODE and DPCLUS) was also compared with our method.

We compared the modularization results on benchmark synthetic LFR networks and on real human PPIN to test the accuracy of modularization after applying the refinement method (Figure 1 and 2). The refinement method sub-optimizes the modularity (quality) of the partition in the pursuit of small modules while maintaining the topological integrity of these modules in order to overcome the resolution limit of modularity-maximizing modular detection algorithms. Present study stresses the fact that the modules with the maximal modularity do not necessarily yield biologically relevant modules in molecular networks. This was evident in the applications to protein-protein interaction networks where a sub-optimal value of the modularity yields modules that are closer to functional modules. We validated this with GO term enrichment as well as with known functionally characterised gene sets.

The improvement in modularization process was observed at different resolution parameters after refinement (Figure 2). To prove that better performance of refined modules is not solely by chance, comparisons are done with the most stable and significant value of resolution parameter (γ=2) that is obtained by comparing the modularization of real networks with multiple random networks of same degree sequence (Figure 3). Refined modules obtained from our method were found to be 4 to 6% more accurate in case of synthetic networks, and in case of protein-protein interaction networks they resulted in better quality modules that were functionally more enriched (Figure 7, 8 and 9) when compared to existing algorithms.

A case study of a supermodule, *X* further substantiated that the algorithm re-modularizes bigger and refinable supermodules into smaller submodules composing of biologically similar proteins (Figure 9). We found that four submodules (A, B, C, D) were rich in specific sets of proteins; *A* having more transporter proteins, *B* more rich in transcriptional regulators, *C* having more transporter, regulator and signalling molecules and *D* moderately rich in signalling molecules given its comparatively smaller size.

Biological processes especially resemble small sub-networks and this new refinement method mines sub-optimal zone of modularity to realize small modules that are functionally more significant. Thus, it produces smaller and refined modules consisting of genes that are functionally more homogeneous and significantly enriched in biological functions.

## Conclusions

The resolution limit of modularization algorithms is a major impediment for the search of computational algorithms that are capable of detecting biologically relevant modules in molecular networks. Our method showed better accuracy in modularization and demonstrated its ability to find smaller modules on synthetic LFR networks mimicking molecular networks and real protein-protein interaction networks. The topological quality and functional significance of the modules realized after applying our method were greatly improved over the existing algorithms. The refinement algorithm developed here is a simple and incremental approach that can extend to other quality-maximizing module detection algorithms to improve the effects of the resolution limit. One could further investigate the convergence properties of our algorithm, the application in overlapping clusters and the effect of the quality loss threshold.

## Methods

### Modularity

#### Modularity as quality function

The *modularity*^5^ is a quantitative measure rendering the quality of the partition of a network into modules. It represents a comparison of edges within a module with the expected number of edges for the nodes in the module for a randomized graph with same size and degree of nodes. Community detection is thus performed by maximizing the modularity. Note that we interchangeably use terms module, community, clusters and sub-graph.

Consider a network *G* with total *L* edges and a partition ℳ = {*m*} of the network. The quality or modularity of its module *m* is given by

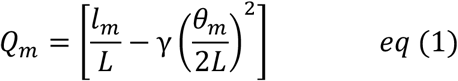

where *l*_*m*_ and *θ*_*m*_ are the number of edges within the module and the total degree of the module, respectively. γ is the resolution parameter with default value of 1. The quality of the modularization ℳ is given by summing the qualities of its modules:

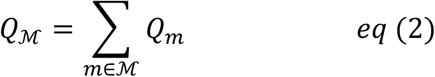

#### Sufficient conditions to be a module

While optimizing the modularity, a subgraph can only be qualified to be a module if total number of intra-module edges is greater than expected by random chance^42,43^, that is, the modularity has to be positive:

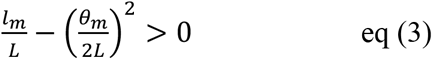

If *l*_*m*_ and 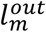 be the number of edges joining nodes within the module *m* and to the rest of the network, respectively, 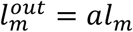 and *a* ≥ 0 is a constant, the degree of the subgraph, 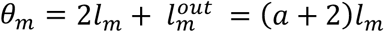. For *a* < 2, the module has a total internal degree 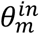 larger than the external degree 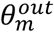 and therefore qualifies to be a module. Therefore, from eq (3) the sufficient conditions for a subgraph to be module are

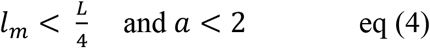

#### Resolution limit of modularity detection

The resolution limit of a module detection algorithm is the smallest size of a module that the algorithm is able to detect. Module detection algorithms using the modularity as the cost function has a resolution limit that depends upon the number of edges between the network modules. From Fortunato and Barthelemy^24^, in a worst case scenario, when intra-module edges (*l*) are balanced with inter-module edges, even larger modules can’t be modularized, and gives maximum scale of resolution limit as

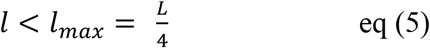

Whereas in a best case scenario when modules are interconnected with only one link, modules that can’t be further modularized will be smaller and gives minimum scale of resolution limit as

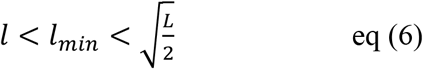

Thus, the modularity has an intrinsic scale of order 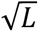 that limits the number and size of modules.

### Module refinement algorithm

#### Sub-optimal values of modularity for functional modules in molecular networks

Modularity optimization, in a best case, cannot detect modules with intra-module edges *l* < *l*_*max*_ and a network can be partitioned to modules with a positive modularity value only if *l* < *l*_*max*_. Thus ideal limits for modularity optimization are [*l*_*min*_, *l*_*max*_]. However, biological networks actually have modules with number of inter-module edges somewhere between two limits, so resolution limit 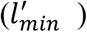 is a bit larger than that of best case. Thus the larger modules will not be detected when 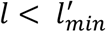 where 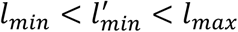.

Therefore, as shown in Figure 10 optimal zone for real biological networks i.e. actual limits for modularity optimization are 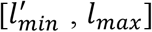. Non-refinable zone is defined as the modules with intra-module edges less than *l*_*min*_ and that cannot be further modularized without disrupting the topological definition of a module. The maximum modularity *Q*_max_ of a real network finds modules with edges having 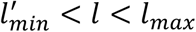. For our purposes, we relax the need for maximum modularity to sub-optimal values *Q*_*ref*_ and re-modularize the bigger modules until we get smaller modules while keeping them topologically relevant untill *l* > *l*_*min*_.

**Figure 10.**
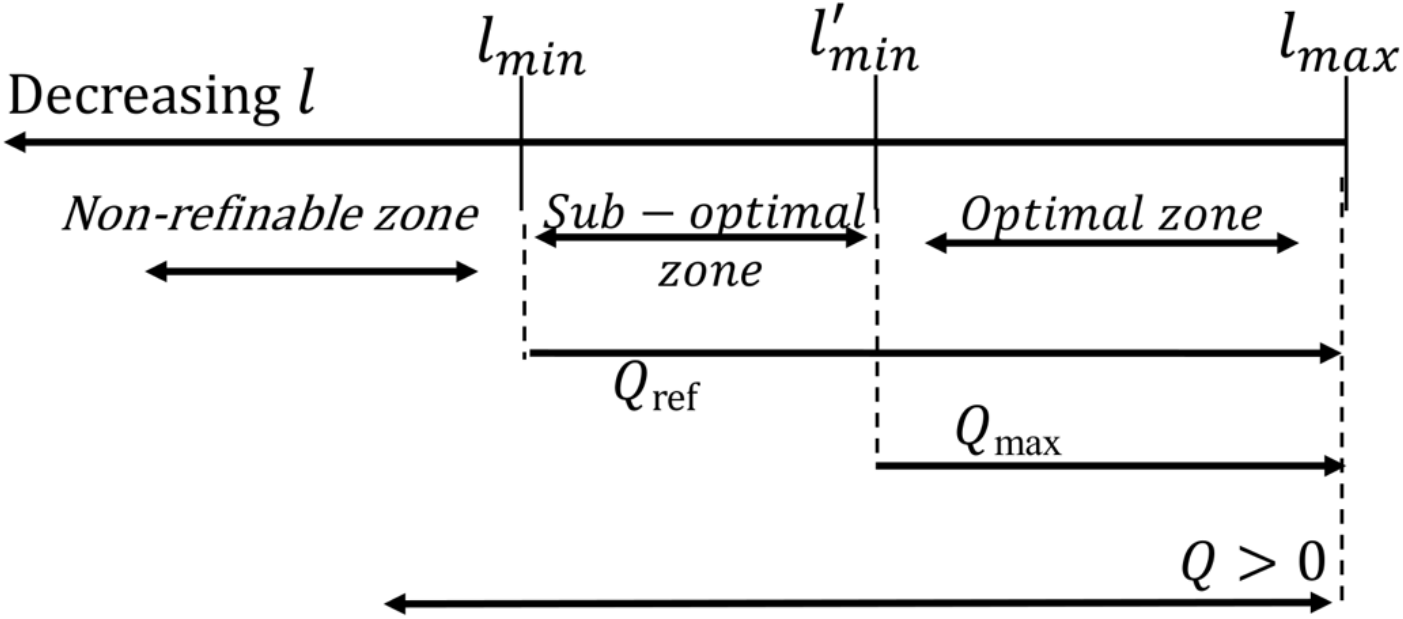
Schematic representation of sub-optimal zone explored for module refinement algorithm.

In this way, our method of module refinement explores the suboptimal zone of modularity values within limits 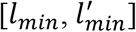. So the sub-optimal modularity *Q*_*ref*_ for a real network is able to find modules such that *l*_*min*_ < *l* < *l*_*max*_.

#### Refinement algorithm

For a given network *G* with *n* nodes and *L* edges, initial modularization of the network is obtained by modularity (*Q*) maximization using existing methods such as Louvain^7^ or Clauset-Newman-Moore Greedy^6^ algorithm. By modularization we mean, partitioning a network into modules. In order to find the smaller modules, larger modules with intra-module edges more than 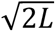 are incrementally modularized such that initial modularization is sub-optimized with the loss of modularity within a user defined specified value (denoted by a threshold value *ρ*).

The steps of the refinement algorithm are:

1. *Initial modularization*: An initial modularization (*M*) obtained by maximizing the modularity *Q* through an existing modularity based algorithm (for example, Louvain) serves as an input for further refinement steps.
2. *Incremental modularization*: According to minimum limit of resolution, modules with edges less than 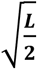 cannot be resolved further (from eq (6)) and we call them *non-refinable* modules. A module is *refinable* if it clubs two or more sub-modules, thus having at least 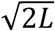 intra-module edges. These larger refinable modules having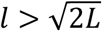 are selected and iteratively re-modularized as individual graphs using the chosen Louvain algorithm. Non-refinable modules are not touched further and added to final modularization (ℳ).
3. *Refinement*: The sub-modules detected in the previous step are filtered using a criteria based on topological constraints: first, the modularity should be positive; and second, the sum of internal degrees 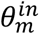 of a module *m* should be greater than the sum of external degrees 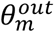 of the module nodes with respect to the rest of the modules. The sub-modules that satisfy these criteria are chosen as it is and those do not satisfy this criteria are grouped in a single subgraph. Both sub-modules are then submitted again for incremental modularization step.
4. *Convergence:* The algorithm converges if there are no more refinable modules left or if the loss of modularity drops below the *threshold loss ρ* (user defined loss of modularity).

Box 1 gives our module refinement algorithm. The notation *M* ← *G* defines a modularization *M* given a graph *G* and *G* ← *M* refers to de-modularization of *M* into a graph *G*.

###### Box 1 Refinement Algorithm

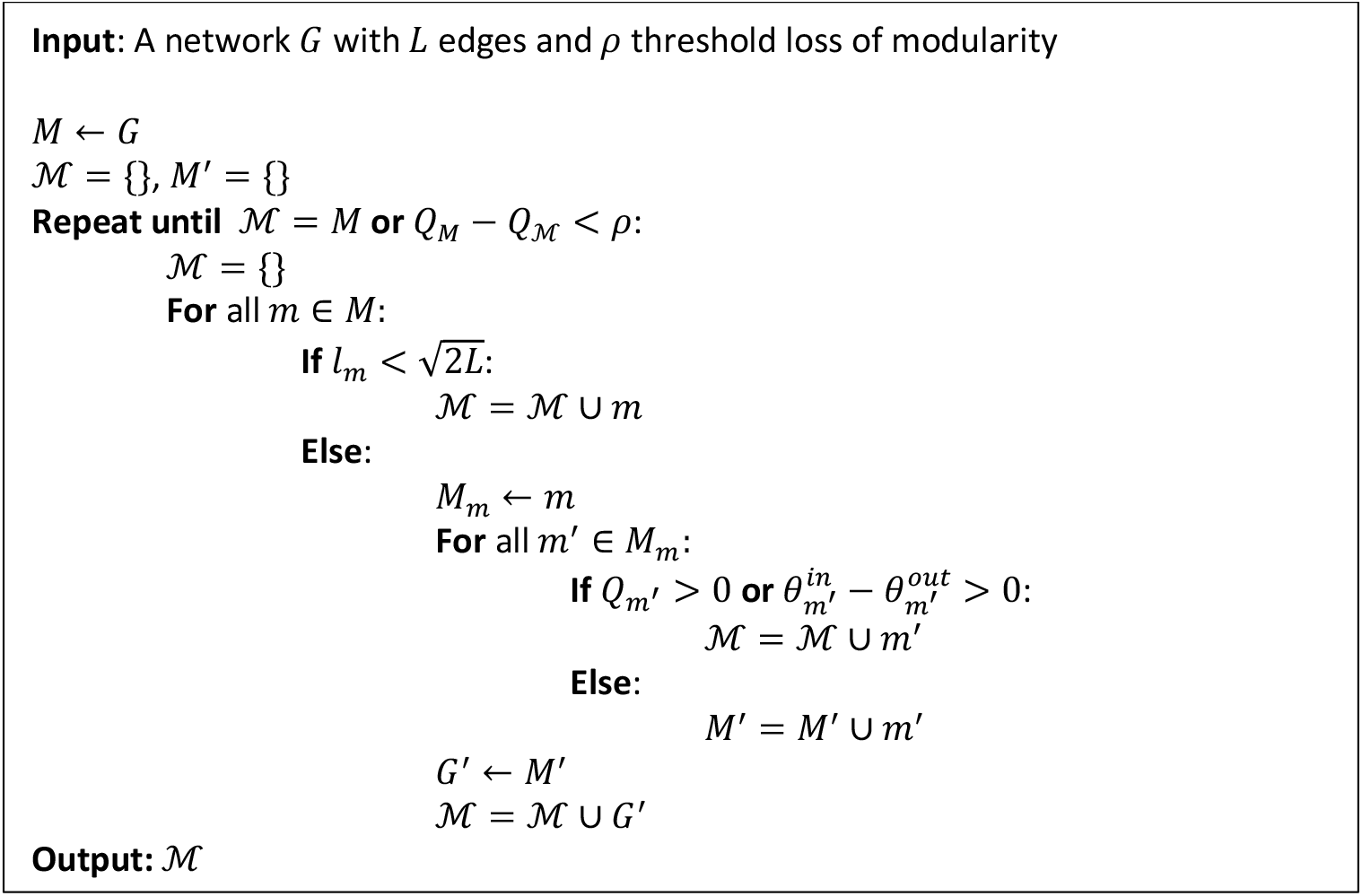

#### Performance evaluation

##### Normalized Mutual Information

For synthetic networks, the accuracy of module detection is estimated using the Normalised Mutual Information (NMI) ^33,44^ between a modularization ℳ = {*m*} (predicted clusters) and the ground truth *T* = {*t*} (real clusters). NMI is also used to estimate stability of partitions obtained over many iterations of module detection.

##### Composite score

In present study, quality of modules is calculated using a composite score which consists of modularity and partition density. Modularity is a global property and measures quality of the partition (eq(2)). Partition density is a local quality measure adapted from^37^ and does not suffer from resolution limit. It is defined as the sum of normalized link density of modules, weighted by the fraction of present edges in the modules. For a network with L edges and modules {m} with intra-module edges *l* and module linked edges *l*_*m*_, partition density

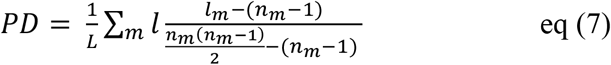

Both Q and PD have values lower than 1. Higher the composite score, better the quality of the partition.

#### Biological Validation

##### Functional enrichment

The biological significance of predicted modules of PPIN was estimated by evaluating the functional enrichment^38^ of the modules, using Gene Ontology (GO) terms^45^ of molecular function (GO-MF). For each module, we focused on the top ten enriched functions for calculating the functional significance of the module. The fraction of genes enriched in significant molecular functions is used as the measure of functional enrichment of the modules.

##### Functional coverage

The functional relevance of modules after refinement was also evaluated by comparing with functionally characterised gene sets given by the Hallmark gene sets^40^ and the KEGG^39^ pathways. The overlap between known functional modules from Hallmark gene sets and KEGG pathways and predicted modules was calculated using Jaccard coefficient. The average overlap between known and predicted modules over various functions gives a measure of functional coverage by the modules of the partition.

#### Other Algorithms

##### Louvain^7^

It finds the modules in a network by optimizing the modularity *Q* (the quality function). Modularization is achieved in two steps. First, all nodes are assigned to individual communities, which are then progressively merged with one of its neighbour’s community that result in best increase in modularity. Secondly, the merged communities are grouped as nodes with edge weight as sum of the edge weights between nodes of each community and first step is repeated. The algorithm is implemented using the python package Community^32^.

Clauset-Newman-Moore Greedy Optimization^6^: It also finds modules in a network by optimizing modularity Q in a similar fashion to Louvain. Only difference is that it merges communities that causes the largest increase in the modularity. The algorithm is implemented using NetworkX^31^.

##### Label Propagation^28^

It assigns communities to a node on the basis of membership of the neighbouring nodes. Initially, each node is assigned a unique community label and then it iteratively updates its community to the most frequent community label among neighbours. The process continues until there is no change in module structure. The algorithm is implemented using NetworkX^31^.

##### Asymptotical Surprise^29^

It optimizes a probability based measure called surprise to find a good partition in the network. The quality function (surprise), *Q* = *mD*(*q*||< *q* >) where *m* is the number of edges, *q* is the fraction of internal edges, < *q* > is the expected fraction of internal edges and *D* is the binary Kullback-Leibler divergence. The quality function is optimised using the Louvain algorithm implemented using the python package Louvain-igraph^46^.

##### MCODE^10^

It detects dense protein complexes in PPIN. It seeds the initial cluster with highest weighted node and then expands these clusters by including neighbour nodes with weights above certain cut-off. The code is implemented to detect non overlapping clusters using the python package Python-graph-clustering^47^.

#### DPCLUS^13^

It also detects protein complexes in PPIN by adding neighbour nodes to the initial cluster that is seeded by the highest weighted node. Clusters are expanded until there is a drop in cluster density and cluster property. The code is implemented to find non overlapping clusters using the python package python-graph-clustering^47^

## Abbreviations

ASY: Asymptotical surprise
G: Greedy optimization algorithm
GO: Gene ontology
GPCR: G-protein coupled receptor
L: Louvain
LFR: Lancichinetti, Fortunato & Radicchi Benchmark networks
LP: Label propagation
MCODE: Molecular Complex Detection
NMI: Normalized mutual information
PD: Partition Density
PPIN: Protein-protein interaction networks
Q: Modularity
TF: Transcription factors

## Declarations

### Ethics approval and consent to participate

Not applicable

### Consent for publication

Not applicable

### Availability of data and material

The source code for algorithm, its usage and data used during the current study is publicly available at https://github.com/ramakaalia/ModuleRefinement.

### Authors’ contributions

RK performed the analysis, interpreted the results and wrote the manuscript. JR supervised the analysis and interpretation of results. All authors have read and approved the final manuscript.

## Acknowledgements

Not applicable

## Additional Files

### Additional file 1

additionalfile.pdf

The file contains supplementary information on dataset pre-processing and analyses supporting the manuscript.

