## Supplementary file for "Refining modules to determine functionally significant clusters in molecular networks"

### Supplementary Information

#### A. Construction of Human Protein-Protein Interaction Networks:

Human protein-protein interaction networks for present study were retrieved from STRING, HPRD, BIOGRID and IMEx consortium databases (DIP, IntAct, MINT, HPIDB, UniProt etc). High confidence physical and functional interactions with score more than 700 were selected from STRING database. All manualy curated physical interactions from HPRD and BioGRID were selected. High score physical interactions with MINT score more than 0.5 were selected from IMEx consortium. Total PPIN dataset contains 78705 interactions and 12022 nodes.

#### B. Normalized Mutual Information:

The overlap between two sets of clusters U and V can be calculated using normalized mutual information as:

$$NMI = \frac{MI(U,V)}{Avg(H(U), H(V)}$$

$$MI(U,V) = \sum_{i=1}^{m} \sum_{j=1}^{n} \frac{n_{ij}}{N} (\log \frac{n_{ij}/N}{a_i b_j/N^2})$$

$$H(U) = -\sum_{i=1}^{m} \frac{a_i}{N} (log \frac{a_i}{N})$$

$$H(V) = -\sum_{i=1}^{n} \frac{b_i}{N} (log \frac{b_i}{N})$$

Where N is total number of nodes, a is number of nodes in clusters of U and b is number of nodes in clusters of V, n is the number of nodes common to clusters of U and V.

#### C. Effect of module refinement:

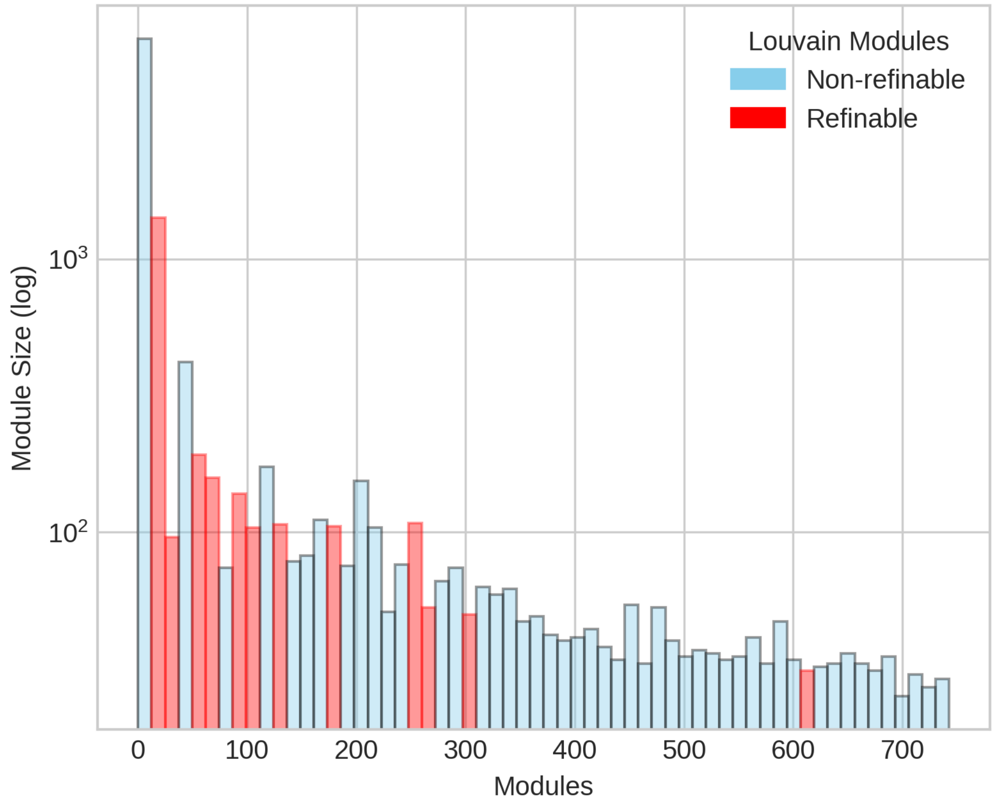


Figure S1. Size distribution of refinable and non-refinable modules obtained from Louvain based modularity optimization, L(γ=2).

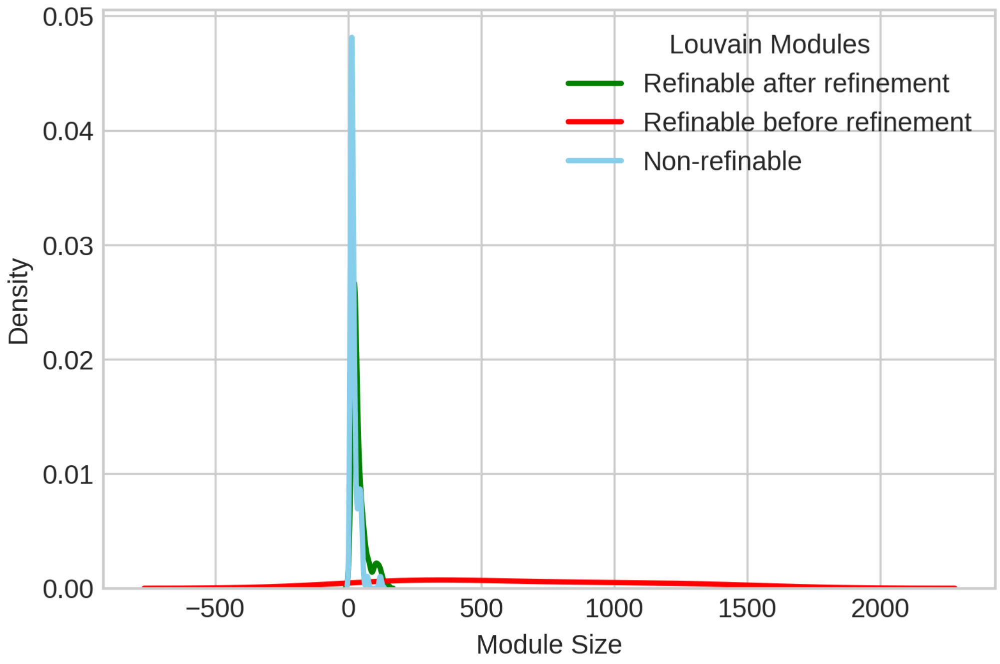


Figure S2. Module size distribution of refinable (from L(γ=2)), non-refinable(from L(γ=2)) and refined (after L(γ=2) +refinement)

### D. Gene list for supermodule X and its submodules A, B, C and D.

The supermodule X is one of the larger refinable modules obtained from partitioning of human protein-protein interaction network using Louvain based modularity optimization (at γ=2) and submodules A, B, C and D are observed from X after refinement.

*Supermodule X:*

6624; 554; 7884; 51384; 9380; 10438; 7415; 6430; 432; 5195; 3762; 3931; 54221; 220082; 6908; 51142; 246744; 84727; 26059; 1544; 64170; 56477; 2185; 100526842; 55226; 140803; 133; 26225; 50514; 51719; 79760; 84557; 2042; 717; 4053; 25927; 140901; 3111; 65259; 9066; 3815; 146279; 27095; 1000; 57187; 6558; 3419; 53832; 6539; 54212; 6881; 57135; 340273; 80781; 57547; 10116; 8735; 64927; 10943; 10906; 347733; 4324; 23020; 84059; 27161; 7162; 27329; 57185; 5890; 10451; 6004; 23623; 115704; 8495; 9340; 241; 93986; 60482; 56940; 55166; 9413; 11334; 79012; 65264; 10956; 1901; 116113; 5329; 54857; 1468;

10868; 6444; 5714; 79188; 4610; 6014; 9581; 23480; 81606; 5609; 10567; 225689; 51645; 10005; 51277; 56157; 11140; 3008; 89953; 123920; 2905; 3266; 5349; 10818; 80153; 1314; 54606; 80204; 832; 381; 51426; 545; 401388; 5864; 3550; 3346; 1399; 54101; 9175; 3755; 9946; 221016; 2646; 5757; 4107; 84639; 11332; 57673; 10963; 2140; 150572; 89853; 55930; 55571; 10273; 5644; 57703; 148738; 84612; 6238; 5584; 9184; 27031; 814; 60592; 2555; 10095; 6374; 1020; 53826; 10477; 54676; 1119; 7917; 22976; 84221; 4920; 3784; 5324; 258010; 3352; 1618; 1718; 56616; 6271; 55898; 2734; 7014; 60673; 23268; 25766; 393; 29968; 4097; 10736; 11066; 10; 4159; 140456; 23389; 8794; 81557; 5338; 5590; 11164; 4515; 6776; 862; 79866; 590; 7678; 29091; 2065; 65983; 10131; 865; 140825; 6450; 80174; 5664; 51654; 81793; 6495; 5300; 54973; 4340; 4541; 338328; 3083; 128710; 23032; 4913; 121549; 10406; 57402; 2794; 3241; 80762; 5927; 54510; 6168; 6658; 7156; 4076; 9188; 100132285; 54870; 147945; 10539; 3920; 3357; 8029; 51168; 55183; 316; 762; 6988; 84191; 1656; 8631; 1620; 23301; 57117; 347240; 8774; 8519; 93129; 11073; 3396; 1186; 9401; 3046; 3218; 221184; 56891; 79576; 6307; 25876; 537; 56904; 7846; 728340; 3037; 118471; 4622; 7421; 26251; 23705; 1938; 5885; 79840; 23042; 50632; 4134; 8892; 8331

*Submodule A:*

5048; 8079; 3337; 22824; 3303; 3304; 55191; 275; 1801; 151194; 51266; 2861; 23640; 192111; 9530; 3309; 65018; 6523; 9992; 89941; 220042; 55585; 3312; 777; 64374; 26098; 10273; 8993; 85300; 573; 114984; 7266; 3306; 11080; 9532; 9451; 3305; 51726; 79982; 1728; 6559; 23746; 5611; 93099; 10294; 10049; 55288; 3757; 10808; 136371; 80022; 9131; 161582; 83983; 3310; 3938; 144245

*Submodule B:*

54788; 27102; 8468; 84129; 11140; 55425; 9931; 2103; 10598; 55631; 26355; 373509; 63943; 23221; 259217; 5681; 728071; 55172; 27330; 93550; 10724; 56950; 11146; 10869; 10910; 4293; 65061; 54802; 3320; 55664; 27101; 5138; 146862; 9049; 164153; 65975; 54970; 5481;

55937; 64427; 79823; 51182; 728642; 2288; 283237; 80222; 10963; 100529239; 2194; 144811; 116835; 79657; 51236; 64754; 29765; 5536; 124540; 150275; 10728; 2289; 85481; 130872; 54851; 6277; 5141; 10953; 741; 6782; 3326; 51447; 57037; 26973; 7268

*Submodule C:*

3154; 1130; 51599; 285527; 984; 10749; 25937; 2810; 5784; 26130; 23367; 23045; 55177; 55119; 23164; 23373; 4291; 23034; 90102; 219771; 5139; 54541; 7538; 5218; 284114; 6546; 55196; 8844; 8711; 56623; 10298; 151; 7352; 57530; 83732; 23387; 137392; 8534; 53616; 23108; 7533; 7531; 1503; 64092; 152; 26064; 7529; 10137; 26031; 9610; 51307

*Submodule D:*

3831; 55898; 22881; 9462; 642273; 6901; 1453; 1452; 474; 56977; 162333; 83891; 60682; 286077; 79856; 81610; 55861; 222584; 1454
